# Between-group conflict participation is associated with lower long-term cortisol in wild vervet monkeys

**DOI:** 10.64898/2026.09.30.755597

**Authors:** Maria Granell Ruiz, Erica van de Waal, Carel P. van Schaik, Redouan Bshary

**Author notes:** Corresponding author: Maria Granell-Ruiz; Pierre-à-Mazel 11, 2000, Neuchâtel, Switzerland.

## Abstract

Male-biased cooperative behaviours that produce public goods (“male services”) are widespread in primates. Recent behavioural studies suggest that male services may function beyond paternal investment, either as displays of competitive ability or as honest signals eliciting female choice. Because these hypotheses make contrasting predictions about the physiological state of service providers, endocrine measures offer a complementary approach for distinguishing between them. We investigated whether long-term testosterone and cortisol concentrations, measured from hair samples, were associated with four male services in wild vervet monkeys: alarm calling, participation in between-group conflicts (BGCs), leading river crossings, and sentinelling. Testosterone and cortisol concentrations increased with male competitive status, measured as a combination of dominance rank and recent rank trajectory, particularly during the mating period, establishing the endocrine profile associated with competition. In contrast, most male services showed few relationships with hormone concentrations: participation in BGCs was associated with lower cortisol concentrations and, to a lesser extent, lower testosterone concentrations. Because BGC participation is the strongest predictor of male mating frequency in this population, these findings indicate that males contributing to BGCs present an endocrine profile distinct from that associated with competitive state and instead support the prediction that high-quality males, irrespective of their dominance rank, can perform costly cooperative behaviours while maintaining relatively low physiological costs. More broadly, our results provide physiological support for reputation-based partner choice as a mechanism contributing to the maintenance of male services in primates.

## INTRODUCTION

Helping behaviours often require individuals to invest time, energy, or exposure to risk to benefit others. Explaining how such behaviours evolved remains a central challenge in evolutionary biology. Helping can evolve through altruism, when actors obtain indirect fitness benefits through kin selection (Hamilton 1964), or through cooperation, when helping yields direct fitness benefits to the actor (Roberts 2005; Bshary and Bergmüller 2008). Identifying these benefits becomes particularly challenging when help produces public goods, where benefits are shared among many group members rather than directed toward specific recipients. One example is the consistent male bias in protective group behaviours observed across many primate species (Carel P. Van Schaik et al. 2022; Granell-Ruiz et al. preprint). Males frequently contribute more than females to predator alarm calling, participation in between-group conflicts (BGCs), leading risky group movements, and sentinelling (DeVore 1963; C. P. Van Schaik and Van Noordwijk 1989; Stephan and Zuberbühler 2016; Baldellou and Peter Henzi 1992; review: Carel P. Van Schaik et al. 2022). These behaviours often involve substantial costs, including increased exposure to predators, energetic expenditure, and elevated risk of injury, while generating benefits for all group members. Because the adult males of a primate group are often not related to the other adults (Pusey and Wolf 1996), and only some have sired offspring, altruism through kin selection is unlikely to explain services directed towards the group as a whole. As a result, male services raise a fundamental question: what direct benefits do males gain by investing in costly protective behaviours that primarily benefit others?

Traditionally, male services have been interpreted under the paternal care hypothesis, whereby males increase their fitness by protecting offspring they have sired. However, this explanation alone appears insufficient to explain the persistence of male services across different reproductive stages in primates (Carel P. Van Schaik et al. 2022). Consequently, alternative hypotheses have proposed that sexual selection may also play an important role, with male services functioning either as honest signals to females through reputation-based partner choice (RBPC) or as displays of competitive ability to rival males (Roberts 1998; Roberts et al. 2021; Dattner et al. 2016; Carel P. Van Schaik et al. 2022). Behavioural analyses in wild vervet monkeys support this broader framework (Granell-Ruiz et al. preprint). Service provision varied across male reproductive careers and with rank: newly immigrated high-ranking males contributed disproportionately to several services, whereas males that had potential offspring in the group invested more consistently across dominance ranks. Yet, dominance rank failed to explain mating frequency in the mating season. Although attaining high dominance rank can provide reproductive advantages, female vervets have considerable leverage over mating (Young et al. 2017; Cheney and Seyfarth 1988; Hemelrijk et al. 2020), and male rank is not consistently associated with mating (Granell-Ruiz et al. preprint) or reproductive success (Minkner et al. 2018). In contrast, while males with higher previous mating frequency invested more in some services, consistent with paternal care, BGC participation and, to a lesser extent, alarm calling predicted male mating frequency more strongly than dominance rank. These patterns may reflect different routes through which males obtain mating benefits, with males potentially differing in their investment in male–male competition for dominance versus services that increase their attractiveness as mating partners.

However, behavioural observations alone cannot determine the relative contribution of these processes because different functions may produce similar patterns of service provision. As these functions may be governed by different underlying biological mechanisms, endocrine measures may help distinguish between processes by revealing whether male services are associated with competitive states or with relatively low physiological costs. Since hormones regulate reproductive effort, competitive interactions, energy allocation, and responses to social challenges, they can be used to characterize these contrasting physiological profiles. Paternal care does not generate a distinctive endocrine prediction, as offspring protection may involve aggression and energetic investment, with hormonal responses depending on the intensity and costs of protection (Muller 2017). This ambiguity is reinforced by previous behavioural data from this population, in which service provision by putative fathers was unrelated to male rank (Granell-Ruiz et al. preprint). We therefore focus on the contrasting physiological predictions of male-male competition and RBPC. Under the male-male competition hypothesis, males providing services are expected to present endocrine profiles associated with investment in acquiring and maintaining dominance. We assess competitive state in relation to a male’s current dominance rank and recent rank dynamics. In contrast, under the RBPC hypothesis, male services are expected to function as honest signals of quality, with male quality reflected by an individual’s ability to repeatedly perform costly cooperative behaviours while maintaining a relatively low physiological burden.

Testosterone and cortisol are particularly informative for distinguishing between these functional explanations. Elevated testosterone concentrations are frequently associated with mate competition, BGC participation, aggression, status acquisition, and reproductive effort (Wingfield et al. 1990; Eisenegger et al. 2011; Hirschenhauser and Oliveira 2006). In primates, these relationships are reflected by higher testosterone concentrations in dominant males and during periods of intense competition for reproductive opportunities (Gesquiere et al. 2011; Muller 2017; Muller and Wrangham 2004a). Testosterone is therefore commonly interpreted as a physiological marker of competitive state. Under the male-male competition hypothesis, males that provide more services should exhibit elevated testosterone concentrations. In contrast, the RBPC hypothesis does not make a specific prediction regarding testosterone.

Cortisol offers complementary information, as it plays a central role in regulating energy mobilisation. Although acute elevations facilitate adaptation to short-term challenges, chronically elevated concentrations can reflect sustained physiological demands (Sapolsky 2005). Elevated cortisol concentrations have also been observed during periods of increased social and reproductive competition, where they may reflect the physiological demands of competing for and maintaining dominant status (Creel 2001; Sapolsky 2005; Gesquiere et al. 2011; Ostner et al. 2008; Thierry et al. 2023). Thus, if male services are associated with this competitive state, service providers may also show elevated cortisol concentrations alongside elevated testosterone. In contrast, if services function as honest signals of quality (RBPC hypothesis), males capable of providing services at relatively low physiological cost may be expected to present lower cortisol concentrations, consistent with the prediction that high-quality individuals can absorb the costs of cooperation more effectively (Zahavi 1975; Leary and Baugh 2020; Moore et al. 2016). Consequently, cortisol may help distinguish whether male services primarily reflect competitive state or function as honest signals of male quality.

Because we were interested in the endocrine profiles associated with consistent service provision rather than acute responses to individual events, hair hormone analysis was particularly well suited for our study. Unlike blood, saliva, urine, or faecal samples, which reflect short-term endocrine activity (from seconds to days), steroid hormones incorporated into growing hair offer an integrated measure of hormonal activity across weeks or months (Davenport et al. 2006; Meyer and Novak 2012; Laudenslager et al. 2012). Hair hormones therefore capture long-term physiological state rather than short-term endocrine fluctuations. This is especially relevant for understanding the evolutionary drivers of male services, as our goal is not to examine the endocrine responses associated with individual service events, but rather the longer-term physiological profiles associated with sustained service provision.

Building directly on our recent behavioural analyses of the same wild vervet monkey population (Granell-Ruiz et al. preprint), we investigated whether long-term testosterone and cortisol concentrations could help distinguish between the proposed mechanisms underlying male services. We first identified the major sources of endocrine variation within the population by examining how testosterone and cortisol concentrations varied with competitive status, social integration, tenure, and season. This allowed us to establish the endocrine profiles associated with competitive state before testing whether males providing services exhibited similar profiles. We then asked whether males contributing disproportionally to four different services—alarm calling, participation in BGCs, leading river crossings, and sentinelling—exhibited endocrine profiles indicative of high competitive state or instead showed profiles more consistent with honest signalling under the RBPC hypothesis. Specifically, if male services primarily reflect competitive state, service providers were expected to exhibit elevated testosterone and cortisol concentrations similar to those associated with dominant, competitively active males. Alternatively, if male services function as honest signals of male quality, service providers were expected to exhibit comparatively lower cortisol concentrations despite engaging in costly cooperative behaviours. By comparing the endocrine profiles associated with competitive state and male service provision, this study aims to provide new insight into the physiological mechanisms underlying male services and the evolutionary processes that maintain them.

## METHODS

### Subjects and darting procedure

The study was conducted at the iNkawu Vervet Project (IVP), located in the Mawana Game Reserve, KwaZulu-Natal, South Africa. Data were collected from habituated wild vervet monkeys (*Chlorocebus pygerythrus*) living in five neighbouring social groups (AK, BD, KB, NH and LT). Vervets live in multimale–multifemale groups characterised by female philopatry and male dispersal.

Hair samples were collected during two darting events, in June 2022 and October 2023. Given the approximately three-month period considered to be represented by the hair samples (see below), the June samples primarily reflect endocrine activity during the preceding mating season, whereas the October samples primarily reflect the preceding winter period and beginning of the baby season. For clarity, we therefore refer to these as the mating-period and winter-period samples, respectively.

Importantly, although these sampling periods occurred in different seasons, both followed the annual mating season, after the majority of matings for that year had already taken place. Hormone measurements were obtained from 18 adult males in June 2022 and 17 adult males in October 2023, with 15 individuals sampled during both darting events (Table S1). Individuals were anaesthetised using Ketamine (2022) or Zoletil (2023) administered via dart gun and temporarily removed from their groups for sample collection. Captures were conducted by a certified veterinarian under approval from the University of KwaZulu-Natal (UKZN) Animal Ethics Committee (AREC/00005320/2023(00020574)), in collaboration with Prof. Colleen Downs, and in accordance with European Research Council and Horizon Europe regulations. Following recovery, monkeys were released back into their social group and monitored for at least two hours which in all cases resulted in normal reintegration. Hair samples were collected by shaving a standardised 4 × 4 cm patch from the lower dorsal region, approximately two finger-widths above the base of the tail. Hair was clipped as close to the skin as possible to ensure complete strands were collected, stored in labelled envelopes at room temperature, and later transported to the University of Neuchâtel, Switzerland, for hormone analysis.

### Hormone analysis

Hormone concentrations from hair samples were quantified using high-performance liquid chromatography (HPLC) following the method developed at the Neuchâtel Platform of Analytical Chemistry (NPAC) (Laubi et al. 2025). Briefly, prior to extraction, hair samples were washed in HPLC-grade isopropanol, gently agitated for 20 s, and dried at 40 °C to remove external contaminants. Samples were then finely cut and homogenised before steroid extraction in methanol containing internal standards. Purification was performed using C18 solid-phase extraction cartridges (Discovery DSC-18, 1 mL, 100 mg), which were conditioned with 100% methanol and equilibrated with 5% methanol prior to sample loading, followed by sequential washes with 5% methanol and hexane, and elution with ethyl acetate. Extracts additionally underwent liquid–liquid extraction before evaporation and reconstitution. Final hormone quantification was carried out using HPLC–MS/MS with appropriate calibration curves. Hormone concentrations are expressed as ng/g hair by correcting extract concentrations for the final reconstitution volume and the mass of hair extracted for each sample. An adaptation from the original protocol consisted of modifications to the hair homogenization and preparation steps to accommodate vervet hair structure, including an initial isopropanol wash, drying, and standardised subsampling prior to grinding (full protocol in OSF, see data availability). Full methodological validation and analytical details are described in (Laubi et al. 2025).

### Behavioural data collection and variable calculation

Behavioural and demographic data were collected year-round by trained field assistants who had previously passed interobserver reliability tests and identification tests performed by the on-site scientific manager.

Competitive state was calculated using two complementary measures of male competitive status: dominance rank and recent rank dynamics. Dominance rank for each male was calculated from dyadic agonistic interactions of adult males within the group occurring during the 12 months preceding the darting date using Elo ratings (EloRating package; Neumann and Kulik 2014), standardised within groups to range from 0 (lowest-ranking male) to 1 (highest-ranking male). To account for recent changes in the hierarchy, we additionally calculated rank trajectory as the difference between Elo scores one week after and one week before the darting date. Using observations from both sides of the darting date allowed us to capture ongoing competitive dynamics that may not yet have been fully reflected in an individual’s Elo rank at the time of sampling. These two measures were combined into a single index of competitive status by summing a male’s standardised dominance rank and rank trajectory. Consequently, males with high dominance rank and an improving rank trajectory received the highest values, whereas males with low rank and/or a declining trajectory received the lowest values. Rank trajectory values ranged from -0.205 to 0.645 (median = 0), with no change in rank for 49% of observations, compared with dominance rank values ranging from 0 to 1 (0.48 ± 0.41). To ensure that our results were not dependent on combining dominance rank and rank trajectory into a composite measure, we repeated all analyses using dominance rank alone. These models produced qualitatively equivalent results and did not alter our main conclusions (Tables S4–S5).

Social integration was quantified using group-standardised social centrality scores derived from grooming interactions recorded during the three months preceding darting using the social indices package (C Neumann 2016). Finally, we calculated male tenure as the time between a male’s immigration into a group and the darting date. Male service provisioning was quantified over the three months preceding hair sampling for four cooperative behaviours: participation in between-group conflicts (BGC), alarm calling, leading river crossings, and sentinelling behaviour. We used this three-month window to approximate the period represented by the hair hormone concentrations, consistent with previous studies using hair in vervets and rhesus macaques (Fairbanks et al. 2011; Laudenslager et al. 2012; Davenport et al. 2006). Alarm calling was calculated from naturally occurring alarm events in which at least one adult male called, excluding encounters involving snakes (Granell-Ruiz et al. preprint). For the present analyses, we included all alarm events rather than restricting analyses to predator alarms only, as predator-only events provided insufficient sample size. BGC participation was quantified from agonistic behaviours directed towards out-group members during encounters between neighbouring groups, while river-crossing leadership was assigned to the first individual leading the group across a broad, open river crossing. For these three event-based behaviours, service provision was calculated as the number of events in which a male participated divided by the number of opportunities available while he was present; events involving unidentified adult males were excluded. Sentinelling was recorded during focal observations and calculated as the proportion of observation time spent visually scanning the surroundings from an elevated, unobstructed position with an active body posture. Summary statistics of observation coverage and behavioural opportunities are provided in Table S1.

### Statistical analyses

All analyses were performed in R version 4.4.2. Hormone concentrations were log-transformed to improve normality, and model assumptions were assessed using the DHARMa package (Hartig and Hartig 2017). For analyses involving leading river crossings, we used a reduced dataset that excluded two males sampled during the mating-period and two sampled during the winter-period, as these individuals belonged to the AK group, whose territory does not contain a river and therefore lacked crossing data.

To investigate whether males providing services presented endocrine profiles consistent with competitive state or with honest signalling under the RBPC hypothesis, we adopted a two-step modelling approach. Given the relatively small sample size, this strategy allowed us to first identify the major sources of endocrine variation within the population and then evaluate whether each male service explained additional variation in hormone concentrations beyond these baseline endocrine profiles. We modelled testosterone and cortisol separately using linear mixed-effects models. First, hormone concentrations were modelled as a function of competitive status, social integration (zCSI), tenure length, darting period (end of the mating season or winter season), and group identity, while including animal identity as a random effect. We then simplified the exploratory models by retaining only predictors that contributed significantly to model fit (without reducing AIC), resulting in a baseline model for each hormone. For testosterone, the final base model included competitive status, darting period and group as predictors, as well as male identity as a random effect. For cortisol, we also included social integration (Table S2).

Second, we evaluated whether male service provision explained additional variation in hormone concentrations while accounting for competitive status and the other predictors retained in the baseline model. We fitted one model for each service (alarm calling, participation in BGCs, leading river crossings, and sentinelling), adding each service variable individually to the corresponding baseline model, to avoid overparameterizing given the sample size. Baseline and service models were compared using likelihood ratio tests (analysis of variance), and differences in AIC were used to assess whether including each service improved model fit (Table S3). To provide an overview of the relationships among hormone concentrations, male service variables, and social predictors, we additionally calculated pairwise Pearson correlations among all continuous variables and visualised them as a correlation heatmap (Fig. S1).

## RESULTS

### Endocrine profiles associated with competitive state

To characterise the endocrine profiles associated with competitive state, we first examined how hormone concentrations varied with season and social factors. Sampling period was a major predictor of both testosterone and cortisol concentrations, highlighting the importance of seasonality in vervet physiology (testosterone: χ^2^(1) = 79.68, p < 0.001, β = 0.73; cortisol: χ^2^(1) = 5.57, p = 0.018, β = 0.32). In both cases, hormone concentrations were higher in the mating period than in the winter period (Fig.1). This pattern was very strong for testosterone, with males sampled after the mating season (4.96 ± 1.77 ng/g) showing almost twice the concentrations of males sampled after winter (2.54 ± 0.54 ng/g). Cortisol showed a similar but weaker seasonal pattern (mating-period: 29.0 ± 15.4 ng/g; winter-period: 22.6 ± 10.3 ng/g).

As predicted, competitive status was another strong predictor of hormone concentrations (Fig.1). Males occupying higher dominance positions and experiencing positive rank trajectories had higher testosterone (χ^2^(1) = 17.91, p < 0.001, β = 0.29) and cortisol concentrations (χ^2^(1) = 9.57, p = 0.002, β= 0.54). For testosterone, this relationship was stronger during the mating-period (estimate = 0.20 ± 0.05, p = 0.001), coinciding with peak reproductive competition, than during the winter-period (estimate = 0.12 ± 0.07, p = 0.06). In contrast, the association between cortisol and competitive status was similar across periods (mating-period: estimate = 0.20 ± 0.11, p = 0.08; winter-period: estimate =0.24 ± 0.12, p = 0.06), suggesting that maintaining a highly competitive position may pose similar physiological demands throughout the year. However, the interaction between period and competitive status was not significant for either hormone (testosterone: χ^2^(1) = 0.88, p = 0.349, β = 0.14; cortisol: χ^2^(1) = 0.09, p = 0.762, β = -0.08). Importantly, repeating the analyses using dominance rank alone instead of competitive status produced equivalent results, indicating that the observed hormonal patterns were not driven by the inclusion of rank trajectory (Tables S4–S5).

Neither hormone was associated with tenure length in the group (testosterone: χ^2^(1) = 0.02, p = 0.882, β = -0.01; cortisol: χ^2^(1) = 0.03, p = 0.872, β = 0.03). Social integration also did not predict testosterone concentrations (χ^2^(1) = 0.80, p = 0.373, β = -0.08), although more socially integrated males tended to have higher cortisol concentrations (χ^2^(1) = 3.00, p = 0.083, β = 0.21). Group membership had no detectable effect on either testosterone (χ^2^(4) = 3.67, p = 0.452) or cortisol concentrations (χ^2^(4) = 1.56, p = 0.816). Results were consistent regardless of whether we considered the full exploratory models or the simplified base models that included only the significant predictors (Table S2).

**Figure 1.**
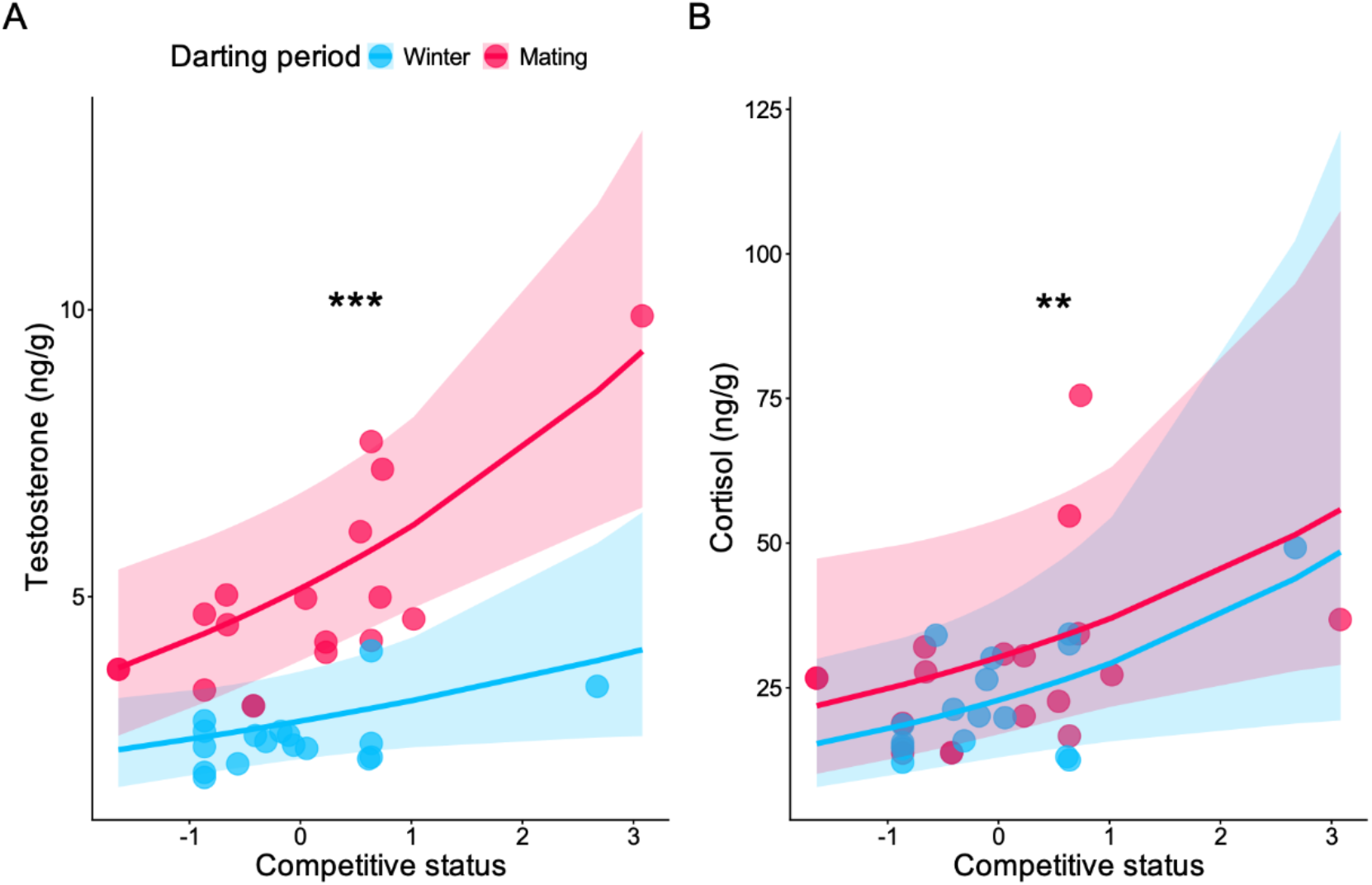
Testosterone and cortisol increase with competitive status. Points show raw testosterone (A) and cortisol (B) concentrations for individual males. Colours indicate darting period (winter-period, red; mating-period, blue). Solid lines show predictions from the exploratory mixed-effects models and shaded areas represent 95% confidence intervals. Testosterone and cortisol increased with competitive status in both periods, although there was a non-significant larger increase for the mating-period in testosterone. Asterisks indicate statistical significance (^**^ p < 0.01, ^***^ p < 0.001).

### Hormone variation and male services

After accounting for the endocrine profiles associated with competitive state, we asked whether male services explained additional variation in hormone concentrations (Fig. 2). BGC participation showed the clearest association with hormone concentrations. Males that participated more frequently in BGCs had significantly lower cortisol concentrations (χ^2^(1) = 5.43, p = 0.020, β = −0.430) and including BGC participation significantly improved model fit (χ^2^(1) = 5.51, p = 0.020). BGC participation showed a similar, although weaker, pattern for testosterone: males that participated more frequently tended to have lower testosterone concentrations (χ^2^(1) = 2.78, p = 0.095, β = −0.161) and adding BGC participation marginally improved model fit (χ^2^(1) = 3.47, p = 0.062). This is the opposite pattern to that observed for male competitive status, where higher-ranking/rank increasing males showed elevated testosterone and cortisol concentrations. Nevertheless, it is important to point out that there does not seem to be a trade-off between competitive state and BGC participation as the correlation coefficient between the two was non-significant but slightly positive (r = 0.23, Fig. S1).

**Figure 2.**
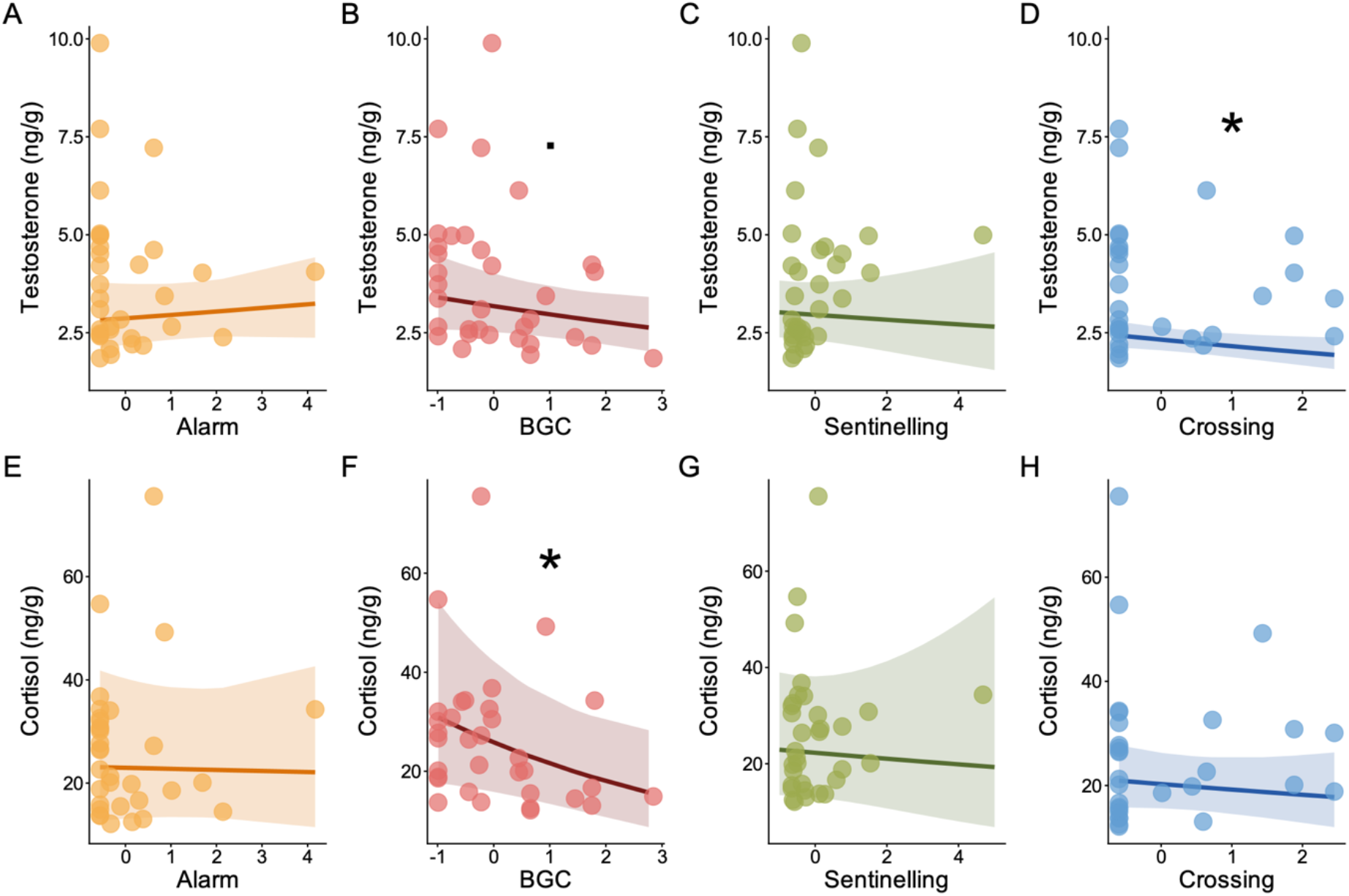
Hormone variation and male services. The upper row shows testosterone (A–D) and the lower row cortisol (E–H) hair hormone concentrations in relation to alarm calling, participation in BGCs, sentinelling and leading river-crossings. Points represent raw hormone values, lines show model predictions, and shaded areas indicate 95% confidence intervals. BGC participation was associated with lower cortisol and testosterone concentrations, while males that led more river crossings showed lower testosterone concentrations. Symbols indicate statistical significance (· p < 0.10, ^*^ p < 0.05).

Similarly, crossing behaviour also showed an association with testosterone: including crossing behaviour marginally improved model fit (χ^2^(1) = 3.48, p = 0.062), and males that led crossings more often had lower testosterone concentrations (χ^2^(1) = 4.73, p = 0.030, β = −0.195). In contrast, alarm calling and sentinelling showed no relationship with either testosterone or cortisol and did not improve model fit (Table S3).

## DISCUSSION

By characterising the endocrine profiles associated with competitive state and comparing them with those associated with male service provision, we investigated whether male services are associated with hormonal patterns more consistent with male-male competition or the reputation-based partner choice (RBPC) hypothesis. As predicted, testosterone and cortisol concentrations were strongly associated with competitive state, with males occupying high-ranking positions and experiencing upward rank trajectories exhibiting elevated concentrations of both hormones. In contrast, participation in male services explained little additional endocrine variation overall. The most interesting exception was BGC participation as this service is most associated with past and future male mating success: males that participated more frequently in BGCs had lower cortisol concentrations and, to a lesser extent, lower testosterone concentrations, an endocrine profile opposite to that associated with competitive state. Leading river crossing was also associated with lower testosterone concentrations, although this pattern was comparatively weak. Together with our previous finding that BGC participation was the strongest predictor of male mating frequency (Granell-Ruiz et al. preprint), these contrasting endocrine profiles provide physiological support for the RBPC hypothesis as a mechanism contributing to the maintenance of male services.

Competitive status is a major determinant of male vervets’ endocrine state. As predicted by the challenge hypothesis, testosterone increased during the mating-period, when reproductive competition is at its highest (Gesquiere et al. 2011; Wingfield et al. 1990). Beyond this seasonal effect, males occupying high-ranking positions or increasing in rank also exhibited elevated testosterone concentrations, suggesting that testosterone reflects not only reproductive competition itself but also the effort required to attain and maintain competitive status (Muller and Wrangham 2004a). Cortisol showed a similar pattern. Because rank in vervets is established and maintained through agonistic interactions, holding a competitive position is likely energetically demanding. While cortisol was also elevated during the mating-period, this effect was much weaker than for testosterone, suggesting that the physiological costs of competition extend beyond periods of reproductive activity (Muller and Wrangham 2004b). Indeed, males at the top of the hierarchy, and particularly those experiencing rank changes, exhibited the highest cortisol concentrations, consistent with evidence that glucocorticoids increase during periods of rank instability and social challenge (Creel 2001; Gesquiere et al. 2011).

This also supports our use of competitive status combining dominance rank with recent rank trajectory. Because hair hormones integrate endocrine activity over an extended period (Behringer and Deschner 2017), incorporating recent rank dynamics may better capture the competitive experience underlying long-term endocrine state than a static measure of rank alone, although analyses using dominance rank alone presented the same overall results. These patterns closely match previous findings in primates and suggest that long-term hormone concentrations reliably capture variation in male competitive state. Together, they reveal a clear endocrine signature of competitive status, with males attaining, maintaining, or improving their position in the hierarchy presenting elevated testosterone and cortisol concentrations.

In contrast, male services did not show the same hormonal profile as competitive status. Most services were unrelated to either testosterone or cortisol, and when associations did emerge, as in BGC and river crossings, they were in the opposite direction. This pattern is difficult to reconcile with the idea that male services are simply another manifestation of male-male competition. If service provision primarily reflected competitive state aimed at increasing mating opportunities, males investing more heavily in these behaviours should exhibit endocrine profiles similar to those associated with high competitive status. Instead, our results suggest that the physiological mechanisms underlying male services differ from those underlying status competition, pointing towards alternative explanations for why males invest in these cooperative behaviours.

In particular, males that participated most frequently in BGC did so while maintaining comparatively low cortisol concentrations. This pattern is consistent with predictions from the RBPC hypothesis, in which cooperative displays function as honest signals of quality because only certain individuals can afford their costs while maintaining good condition (Roberts et al. 2021; Zahavi 1975; Gintis et al.2001; Roberts 1998). Chronically elevated glucocorticoid concentrations are generally associated with sustained energetic demands and can impair immune function, growth, and reproductive performance (Sapolsky 2005; Romero et al. 2009). Thus, rather than indicating that participation in BGC is cost-free, lower cortisol concentrations may suggest that some males are better able to absorb these costs without experiencing prolonged physiological strain. Interestingly, participation in BGCs also tended to be associated with lower testosterone concentrations, a pattern that further argues against BGC participation being primarily driven by male-male competition, while remaining fully consistent with the RBPC hypothesis. In this sense, BGC participation may act as a particularly informative signal of male quality: not because high-quality males avoid costly behaviours, but because they can perform them while maintaining physiological homeostasis (Husak and Moore 2008). Because hair cortisol concentrations show moderate heritability in vervet monkeys (h^2^ = 0.31; Fairbanks et al. 2011), it is possible that variation in physiological resilience has a genetic component that females could assess through observing males participating in BGC. If so, females preferring males that can afford to or are willing to contribute to BGC while maintaining low cortisol concentrations may obtain additional indirect genetic benefits for their offspring (Zahavi 1975; Husak and Moore 2008). Notably, BGC participation is also the strongest predictor of the number of matings obtained by males in this population, outperforming rank, tenure and all other male services (Granell-Ruiz et al. preprint), while females actively monitor male participation during intergroup encounters (Arseneau-Robar et al.2016). Taken together, these findings support the hypothesis that participation in BGCs functions as an honest signal of male quality and are consistent with the idea that female choice has favoured male-biased investment in BGC participation through RBPC.

An important question raised by these findings is why males that are able to _provide_ potentially costly services at relatively low physiological burden are not necessarily those occupying the highest dominance positions. Importantly, this contrast does not suggest a trade-off between the two, as competitive status and BGC participation were weakly positively correlated (r = 0.23; Fig. S1). One possibility is that dominance and service provision represent partly distinct routes through which males obtain mating benefits. High rank may provide advantages through male–male competition, whereas service provision may increase male attractiveness through female choice. In vervets, where females have considerable leverage over mating (Young et al. 2017; Hemelrijk et al. 2020; Cheney and Seyfarth 1988), and dominance rank is not tightly coupled to reproductive success (Minkner et al. 2018), males may differ in how they allocate effort between these routes. Under this interpretation, male quality is not synonymous with dominance rank: quality may instead influence a male’s capacity to invest successfully in either, or potentially both, competitive status and female-attractive services. Such investment decisions may also change across a male’s career as his competitive position and opportunities change (Hill et al. 2026; Van Noordwijk and Van Schaik 2004). Our data cannot determine whether males actively pursue alternative strategies or whether investment changes across their tenure and resolving this will ultimately require direct measures of reproductive success and how these relationships change across male reproductive careers. Nevertheless, this provides a potential explanation for the contrasting physiological profiles associated with competitive status and BGC participation.

Unlike BGC participation, the remaining services provided little endocrine evidence supporting any particular functional hypothesis. Alarm calling, despite being linked to higher mating frequency in behavioural analyses (Granell-Ruiz et al. preprint), was unrelated to either testosterone or cortisol concentrations. One possibility is that alarm calling is in general less physiologically demanding than BGC participation and/or that high-risk alarm events occur too infrequently to generate detectable differences in long-term endocrine state. Indeed, because predator-specific alarm events were relatively rare, we broadened the definition of alarm calling for the present analyses, potentially reducing our ability to detect endocrine patterns specifically associated with high-risk alarm events. Similarly, sentinelling showed no hormonal associations. Given that low-ranking males contribute disproportionately to sentinelling behaviour (Granell-Ruiz et al. preprint), it could reflect exposure to social stress arising from the presence of dominant individuals. However, its lack of association with cortisol suggests that sentinelling does not impose a strong long-term physiological burden, particularly compared with maintaining a high competitive position (Creel 2001; Gesquiere et al.2011). River-crossing leadership remains more difficult to interpret. Males leading crossings tended to present lower testosterone concentrations, this pattern contrasts with our behavioural interpretation of crossing leadership as potentially linked to male–male competition (Granell-Ruiz et al. preprint), which would instead predict an endocrine profile similar to that associated with competitive state. However, this effect was comparatively weak, was not accompanied by a corresponding cortisol effect, and should therefore be interpreted cautiously. Importantly, our dataset does not include any newly immigrated top-ranking males, which have been shown to predominantly lead river crossings (Tankink et al. 2026; Granell-Ruiz et al. preprint). Consequently, the present data may not capture the endocrine profile of the males that typically perform this behaviour, and further work with a more representative dataset is needed.

## Conclusion

Our results demonstrate that BGC participation is associated with an endocrine profile distinct from that associated with male competitive status and consistent with predictions from the RBPC hypothesis. While testosterone and cortisol concentrations increased with rank and recent upward changes in rank, participation in BGCs was associated with lower cortisol concentrations and tended to be associated with lower testosterone concentrations. These findings suggest that BGC participation is not simply an expression of male–male competition but may instead signal a male’s ability to provide a costly cooperative service while maintaining relatively low physiological costs. More broadly, none of the other male services showed the endocrine profile expected under male–male competition. Although our sample size was limited by the challenges of collecting hair hormone data from wild primates, and hair hormones cannot distinguish long-term endocrine state from acute responses to specific events, future studies combining longitudinal endocrine measures with genetic paternity data will help clarify the links between male quality, service provision, and reproductive success. Together with our previous finding that BGC participation strongly predicts male mating frequency, these results raise the possibility that males may pursue different, potentially complementary routes to reproductive success, through investment in competitive status or in services favoured through female choice. By linking long-term endocrine state to cooperative behaviour in a natural population, this study provides a novel physiological perspective on the evolutionary mechanisms that maintain male services in primates.

## DECLARATION OF GENERATIVE AI AND AI-ASSISTED TECHNOLOGIES IN THE MANUSCRIPT PREPARATION PROCESS

During the preparation of this work, the authors used ChatGPT (OpenAI) to assist with language editing and improving the clarity and structure of the manuscript. The authors reviewed and edited all AI-assisted content and take full responsibility for the content of the published article.

## ACKNOWLEDGMENTS

We are grateful to everyone who makes the long-term work of the iNkawu Vervet Project possible and for their dedication to monitoring the study population through thick and thin. We especially thank the field team for their continued hard work (including plenty of digging), their invaluable assistance during captures, and the care with which they helped us handle sedated individuals and navigate the challenges of darting. In particular, we thank Michael Henshall, Nokubonga Dlamini, Josefien Tankink, Siboniso Thela, and Zonke Mbutho for managing the field site and providing invaluable assistance in the field, and Loïc Brun, Michel Halbwax, Lukas Schad and Charlotte Canteloup for their contributions to the darting work. We are also grateful to Benjamin Laubi, Philippe Vullioud, Gaëtan Glauser, and Sylvie Guinchard for their guidance, training, and assistance with hormone analyses. Finally, we thank the van der Walt family for allowing us to conduct research on their reserve and Ezemvelo KZN Wildlife for granting permission to conduct research on vervet monkeys on Mawana, and the University of KwaZulu-Natal (UKZN) Animal Ethics Committee for ethical approval of the capture procedures (AREC/00005320/2023(00020574)).

## CREDIT AUTHORSHIP CONTRIBUTION STATEMENT

Maria Granell Ruiz: Conceptualization, Data curation, Formal analysis, Investigation, Methodology, Software, Validation, Visualization, Writing – original draft, Writing – review and editing. Erica van de Waal: Funding acquisition, Methodology, Project administration, Resources, Writing – review and editing. Carel P. van Schaik: Conceptualization, Funding acquisition, Supervision, Writing – review and editing.

Redouan Bshary: Conceptualization, Funding acquisition, Supervision, Writing – review and editing.

## DATA AVAILABILITY

All data and code required to reproduce the analyses are publicly available. Analysis code is available on GitHub (https://github.com/mariagranell/MS_hair_hormones), while the associated data and the protocol used for hair hormone analysis are archived on OSF (https://osf.io/4mbgd/overview).

## REFERENCES

Arseneau-Robar T. Jean Marie, Anouk Lisa Taucher, Eliane Müller, Carel Van Schaik, Redouan Bshary, and Erik P. Willems. 2016. ‘Female Monkeys Use Both the Carrot and the Stick to Promote Male Participation in Intergroup Fights’. Proc R Soc B 283 (1843): 20161817. 10.1098/rspb.2016.1817.

Baldellou, Maribel, and S. Peter Henzi. 1992. ‘Vigilance, Predator Detection and the Presence of Supernumerary Males in Vervet Monkey Troops’. Animal Behaviour 43 (3): 451–61. 10.1016/s0003-3472(05)80104-6.

Behringer, Verena, and Tobias Deschner. 2017. ‘Non-Invasive Monitoring of Physiological Markers in Primates’. Hormones and Behavior 91 (May): 3–18. 10.1016/j.yhbeh.2017.02.001.

Bshary, R., and R. Bergmüller. 2008. ‘Distinguishing Four Fundamental Approaches to the Evolution of Helping’. Journal of Evolutionary Biology 21 (2): 405–20. 10.1111/j.1420-9101.2007.01482.x.

Cheney, Dorothy L., and Robert M. Seyfarth. 1988. ‘Assessment of Meaning and the Detection of Unreliable Signals by Vervet Monkeys’. Animal Behaviour 36 (2): 477–86. 10.1016/S0003-3472(88)80018-6.

Creel, Scott. 2001. ‘Social Dominance and Stress Hormones’. Trends in Ecology & Evolution 16 (9): 491–97. 10.1016/S0169-5347(01)02227-3.

Dattner, Arnon, Amotz Zahavi, and Avishag Zahavi. 2016. ‘Competition over Guarding in the Arabian Babbler (Turdoides Squamiceps), a Cooperative Breeder’. F1000Research 4: 618.

Davenport, Matthew D., Stefan Tiefenbacher, Corrine K. Lutz, Melinda A. Novak, and Jerrold S. Meyer. 2006. ‘Analysis of Endogenous Cortisol Concentrations in the Hair of Rhesus Macaques’. General and Comparative Endocrinology 147 (3): 255–61. 10.1016/j.ygcen.2006.01.005.

DeVore, Irven. 1963. ‘A Comparison of the Ecology and Behavior of Monkeys and Apes’. In Classification and Human Evolution. Viking Fund Publications in Anthropology.

Eisenegger, Christoph, Johannes Haushofer, and Ernst Fehr. 2011. ‘The Role of Testosterone in Social Interaction’. Trends in Cognitive Sciences 15 (6): 263–71. 10.1016/j.tics.2011.04.008.

Fairbanks, Lynn A., Matthew J. Jorgensen, Julia N. Bailey, Sherry E. Breidenthal, Rachel Grzywa, and Mark L. Laudenslager. 2011. ‘Heritability and Genetic Correlation of Hair Cortisol in Vervet Monkeys in Low and Higher Stress Environments’. Psychoneuroendocrinology 36 (8): 1201–8. 10.1016/j.psyneuen.2011.02.013.

Gesquiere, Laurence R., Niki H. Learn, M. Carolina M. Simao, Patrick O. Onyango, Susan C. Alberts, and Jeanne Altmann. 2011. ‘Life at the Top: Rank and Stress in Wild Male Baboons’. Science 333 (6040): 357–60. 10.1126/science.1207120.

Gintis, Herbert, Eric Alden Smith, and Samuel Bowles. 2001. ‘Costly Signaling and Cooperation’. Journal of Theoretical Biology 213 (1): 103–19. 10.1006/jtbi.2001.2406.

Granell-Ruiz, Maria, Josefien A. Tankink, Erica van de Waal, and Carel P. van Schaik. preprint. Beyond Paternal Care: Career Stage and Reproductive Opportunities Shape Male Services in Vervet Monkeys. 10.64898/2026.06.22.733760.

Hamilton, William D. 1964. ‘The Genetical Evolution of Social Behaviour. II’. Journal of Theoretical Biology 7 (1): 17–52.

Hartig, Florian, and Maintainer Florian Hartig. 2017. ‘Package “Dharma”‘. R Package 531 (532):435. 10.32614/CRAN.package.DHARMa.

Hemelrijk, Charlotte Korinna, Matthias Wubs, Gerrit Gort, Jennifer Botting, and Erica Van De Waal. 2020. ‘Dynamics of Intersexual Dominance and Adult Sex-Ratio in Wild Vervet Monkeys’. Frontiers in Psychology 11 (May). 10.3389/fpsyg.2020.00839.

Hill, Alice, Suchinda Malaivijitnond, Suthirote Meesawat, Oliver Schülke, and Julia Ostner. 2026. ‘Male Careers in Wild Assamese Macaques’. Behavioral Ecology 37 (5): arag085.

Hirschenhauser, Katharina, and Rui F. Oliveira. 2006. ‘Social Modulation of Androgens in Male Vertebrates: Meta-Analyses of the Challenge Hypothesis’. Animal Behaviour 71 (2): 265–77. 10.1016/j.anbehav.2005.04.014.

Husak, Jerry F., and Ignacio T. Moore. 2008. ‘Stress Hormones and Mate Choice’. Trends in Ecology & Evolution 23 (10): 532–34. 10.1016/j.tree.2008.06.007.

Laubi, Benjamin, Gaëtan Glauser, Erik P. Willems, Judith Burkart, and Carel Van Schaik. 2025. ‘Insights into Hair Steroid Dynamics in Male and Female Common Marmosets (Callithrix Jacchus) Using a Novel Hplc-Ms/Ms Assay’. Preprint, SSRN. 10.2139/ssrn.5124513.

Laudenslager, Mark L., Matthew J. Jorgensen, and Lynn A. Fairbanks. 2012. ‘Developmental Patterns of Hair Cortisol in Male and Female Nonhuman Primates: Lower Hair Cortisol Levels in Vervet Males Emerge at Puberty’. Psychoneuroendocrinology 37 (10): 1736–39. 10.1016/j.psyneuen.2012.03.015.

Leary, Christopher J., and Alexander T. Baugh. 2020. ‘Glucocorticoids, Male Sexual Signals, and Mate Choice by Females: Implications for Sexual Selection’. General and Comparative Endocrinology 288 (March): 113354. 10.1016/j.ygcen.2019.113354.

Meyer, Jerrold S., and Melinda A. Novak. 2012. ‘Minireview: Hair Cortisol: A Novel Biomarker of Hypothalamic-Pituitary-Adrenocortical Activity’. Endocrinology 153 (9): 4120–27. 10.1210/en.2012-1226.

Minkner, Mirjam M. I., Christopher Young, Federica Amici, et al. 2018. ‘Assessment of Male Reproductive Skew via Highly Polymorphic STR Markers in Wild Vervet Monkeys, Chlorocebus Pygerythrus’. Journal of Heredity, ahead of print, September. 10.1093/jhered/esy048.

Moore, Fhionna R., David M. Shuker, and Liam Dougherty. 2016. ‘Stress and Sexual Signaling: A Systematic Review and Meta-Analysis’. Behavioral Ecology 27 (2): 363–71. 10.1093/beheco/arv195.

Muller, Martin N. 2017. ‘Testosterone and Reproductive Effort in Male Primates’. Hormones and Behavior 91 (May): 36–51. 10.1016/j.yhbeh.2016.09.001.

Muller, Martin N., and Richard W. Wrangham. 2004a. ‘Dominance, Aggression and Testosterone in Wild Chimpanzees: A Test of the “Challenge Hypothesis”‘. Animal Behaviour 67 (1): 113–23. 10.1016/j.anbehav.2003.03.013.

Muller, Martin N., and Richard W. Wrangham. 2004b. ‘Dominance, Cortisol and Stress in Wild Chimpanzees (Pan Troglodytes Schweinfurthii)’. Behavioral Ecology and Sociobiology 55 (4): 332–40. 10.1007/s00265-003-0713-1.

Neumann, C. 2016. ‘Socialindices: Social and Association Indices’. Unpublished R Package (v. 0. 46–7).

Neumann, Christof, and Lars Kulik. 2014. ‘EloRating: Animal Dominance Hierarchies by Elo Rating’. CRAN: Contributed Packages.

Ostner, Julia, Peter Kappeler, and Michael Heistermann. 2008. ‘Androgen and Glucocorticoid Levels Reflect Seasonally Occurring Social Challenges in Male Redfronted Lemurs (Eulemur Fulvus Rufus)’. Behavioral Ecology and Sociobiology 62 (4): 627–38.

Pusey, Anne, and Marisa Wolf. 1996. ‘Inbreeding Avoidance in Animals’. Trends in Ecology & Evolution 11 (5): 201–6.

Roberts, Gilbert. 1998. ‘Competitive Altruism: From Reciprocity to the Handicap Principle’. Proceedings of the Royal Society of London. Series B: Biological Sciences 265 (1394): 427–31.

Roberts, Gilbert. 2005. ‘Cooperation through Interdependence’. Animal Behaviour 70 (4): 901–8.

Roberts, Gilbert, Nichola Raihani, Redouan Bshary, et al. 2021. ‘The Benefits of Being Seen to Help Others: Indirect Reciprocity and Reputation-Based Partner Choice’. Philosophical Transactions of the Royal Society B: Biological Sciences 376 (1838): 20200290. 10.1098/rstb.2020.0290.

Romero, L. Michael, Molly J. Dickens, and Nicole E. Cyr. 2009. ‘The Reactive Scope Model — A New Model Integrating Homeostasis, Allostasis, and Stress’. Hormones and Behavior 55 (3): 375–89. 10.1016/j.yhbeh.2008.12.009.

Sapolsky, Robert M. 2005. ‘The Influence of Social Hierarchy on Primate Health’. Science 308 (5722): 648–52. 10.1126/science.1106477.

Stephan, Claudia, and Klaus Zuberbühler. 2016. ‘Persistent Females and Compliant Males Coordinate Alarm Calling in Diana Monkeys’. Current Biology 26 (21): 2907–12. 10.1016/j.cub.2016.08.033.

Tankink, Josefien A., Erica van de Waal, Redouan Bshary, and Carel P. van Schaik. 2026. ‘Leadership Under Risk: Male Vervet Monkeys’(Chlorocebus Pygerythrus) Roles in Group Progression Across Potentially High-Risk Terrain’. Ethology, ahead of print. 10.1111/eth.70085.

Thierry, Bernard, Nancy Rebout, and Michael Heistermann. 2023. ‘Hormonal Responses to Mating Competition in Male Tonkean Macaques’. Hormones and Behavior 154 (August): 105395. 10.1016/j.yhbeh.2023.105395.

Van Noordwijk, Maria A., and Carel P. Van Schaik. 2004. ‘Sexual Selection and the Careers of Primate Males: Paternity Concentration, Dominance-Acquisition Tactics and Transfer Decisions’. In Sexual Selection in Primates, 1st edn,edited by Peter M. Kappeler and Carel P. Van Schaik. Cambridge University Press. 10.1017/CBO9780511542459.014.

Van Schaik, C. P., and M. A. Van Noordwijk. 1989. ‘The Special Role of Male Cebus Monkeys in Predation Avoidance and Its Effect on Group Composition’. Behav Ecol and Sociobiology 24 (5): 265–76. 10.1007/bf00290902.

Van Schaik, Carel P., Redouan Bshary, Gretchen Wagner, and Filipe Cunha. 2022. ‘Male Anti-predation Services in Primates as Costly Signalling? A Comparative Analysis and Review’. Ethology 128 (1): 1–14. 10.1111/eth.13233.

Wingfield, John C., Robert E. Hegner, Alfred M. Dufty, and Gregory F. Ball. 1990. ‘The “Challenge Hypothesis”: Theoretical Implications for Patterns of Testosterone Secretion, Mating Systems, and Breeding Strategies’. The American Naturalist 136 (6): 829–46. 10.1086/285134.

Young, Christopher, Richard McFarland, Louise Barrett, and S. Peter Henzi. 2017. ‘Formidable Females and the Power Trajectories of Socially Integrated Male Vervet Monkeys’. Animal Behaviour 125 (March): 61–67. 10.1016/j.anbehav.2017.01.006.

Zahavi, Amotz. 1975. ‘Mate Selection—A Selection for a Handicap’. Journal of Theoretical Biology 53 (1): 205–14. 10.1016/0022-5193(75)90111-3.

