## Supplementary materials for "Between-group conflict participation is associated with lower long-term cortisol in wild vervet monkeys"

APPENDIX

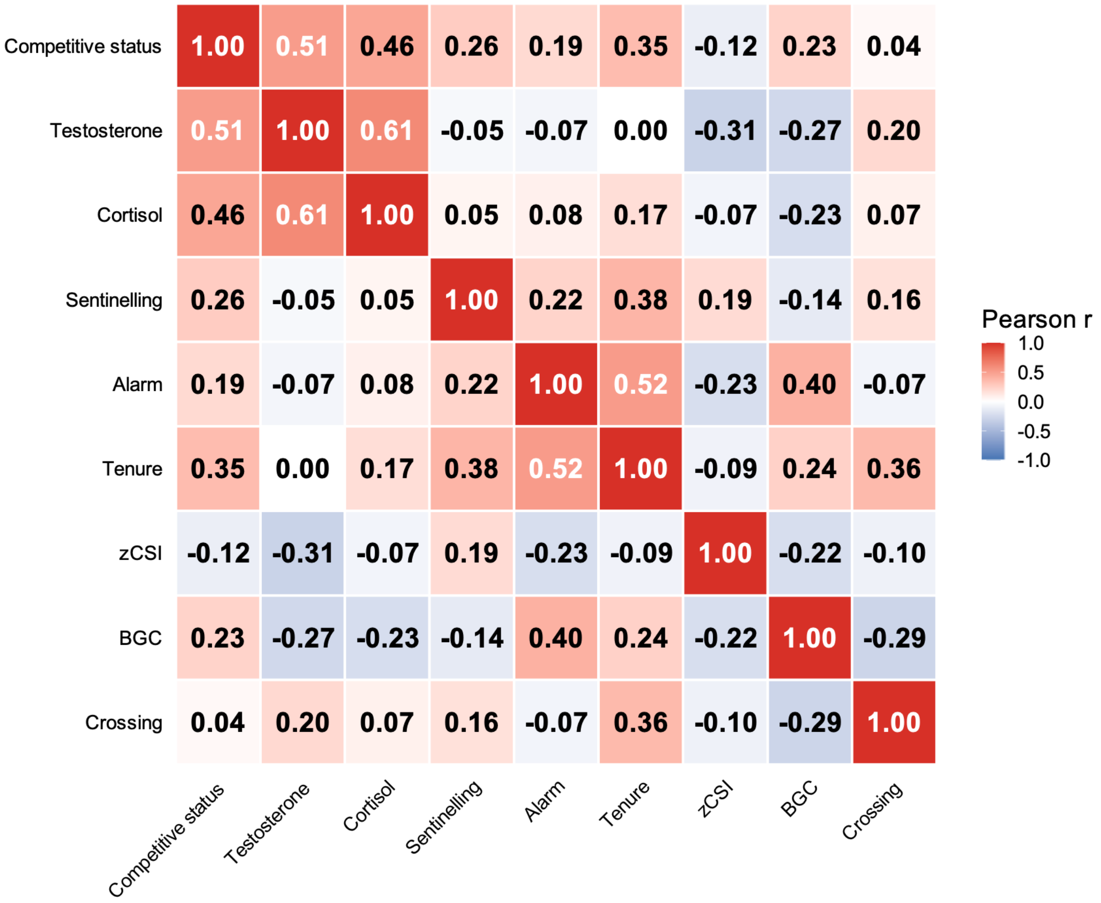


Figure S1. **Correlation matrix of hormone concentrations and male service behaviours.** Pearson correlation coefficients among testosterone and cortisol concentrations (ng/g), four male service behaviours (proportion of alarm calling, between-group encounter participation, river crossing leadership, and sentinelling), social integration (zCSI), tenure and competitive status (rank and rank trajectory). Cell colours indicate the magnitude and direction of the correlation, and values within cells show Pearson's r. Positive correlations are shown in red and negative correlations in blue.

| Season | Group | N | Days Present | Sampling Effort (%) | Focal (min) | Alarms | BGC | Crossings |
|---|---|---|---|---|---|---|---|---|
| Winter | AK | 2 | 59.5 ± 0.7 | 64.0 ± 0.8 | 165.8 ± 11.3 | 7.5 ± 0.7 | 11.0 ± 0.0 | - |
| | BD | 7 | 69.1 ± 1.2 | 74.3 ± 1.3 | 112.5 ± 56.2 | 40.6 ± 1.1 | 7.4 ± 0.8 | 15.4 ± 0.8 |
| | KB | 2 | 56.5 ± 2.1 | 60.8 ± 2.3 | 198.8 ± 56.0 | 19.0 ± 1.4 | 6.0 ± 0.0 | 9.5 ± 0.7 |
| | LT | 2 | 44.5 ± 0.7 | 29.4 ± 0.2 | 36.0 ± 1.6 | 2.0 ± 0.0 | 4.0 ± 0.0 | 2.0 ± 0.0 |
| | NH | 4 | 59.5 ± 0.6 | 64.0 ± 0.6 | 122.7 ± 53.5 | 40.8 ± 2.5 | 20.8 ± 0.5 | 6.2 ± 1.0 |
| Mating | AK | 3 | 34.3 ± 27.2 | 52.5 ± 2.4 | 16.7 ± 4.0 | 5.0 ± 0.0 | 8.0 ± 0.0 | - |
| | BD | 6 | 52.7 ± 13.7 | 61.3 ± 7.5 | 37.7 ± 23.2 | 21.2 ± 0.4 | 15.3 ± 2.2 | 7.5 ± 2.9 |
| | KB | 1 | 37.0 | 39.8 | 12.1 | 6.0 | 5.0 | 1.0 |
| | LT | 1 | 46.0 | 37.4 | 34.8 | 4.0 | 5.0 | 2.0 |
| | NH | 6 | 43.5 ± 17.1 | 61.9 ± 17.1 | 34.5 ± 17.9 | 10.7 ± 7.9 | 9.5 ± 4.6 | 2.5 ± 2.3 |

Table S1. Summary of sample sizes, observation coverage, and behavioural opportunities available to males included in the hormone analyses during the three-month period preceding the darting of each male. N indicates the number of males sampled per group. Values are presented as mean ± SD. Sampling effort (%) was calculated as the proportion of days an individual was present in the group during which behavioural observations were conducted. Mean focal observation time is reported in minutes. Alarms, BGCs, and crossings represent the mean number of behavioural opportunities available to males during the corresponding three-month sampling window. The AK territory does not contain a river crossing site and was therefore excluded from crossing analyses.

MODEL OUTPUTS

| | | Model | n | Exploratory vs Base | Competitive status | Darting Period | Competitive status × Season | Tenure | zCSI | Group |
|---|---|---|---|---|---|---|---|---|---|---|
| Testosterone | Full data | Exploratory | 35 | $\chi^2(3)=2.40$, p=0.494 | **$\chi^2(1)=17.91$, p<0.001 [β=0.286]** | **$\chi^2(1)=79.68$, p<0.001 [β=0.730]** | $\chi^2(1)=0.88$, p=0.349 [β=0.138] | $\chi^2(1)=0.02$, p=0.882 [β=-0.014] | $\chi^2(1)=0.80$, p=0.373 [β=-0.082] | $\chi^2(4)=3.67$, p=0.452 |
| | | Base | 35 | | **$\chi^2(1)=19.34$, p<0.001 [β=0.378]** | **$\chi^2(1)=102.13$, p<0.001 [β=0.756]** | — | — | — | $\chi^2(4)=5.47$, p=0.242 |
| | Crossing subset | Exploratory | 33 | $\chi^2(3)=1.53$, p=0.676 | **$\chi^2(1)=12.31$, p<0.001 [β=0.288]** | **$\chi^2(1)=75.57$, p<0.001 [β=0.849]** | $\chi^2(1)=0.74$, p=0.388 [β=0.125] | $\chi^2(1)=0.24$, p=0.623 [β=-0.057] | $\chi^2(1)=0.08$, p=0.776 [β=-0.032] | $\chi^2(4)=5.03$, p=0.285 |
| | | Base | 33 | | **$\chi^2(1)=15.16$, p<0.001 [β=0.350]** | **$\chi^2(1)=100.61$, p<0.001 [β=0.859]** | — | — | — | $\chi^2(4)=6.03$, p=0.197 |
| Cortisol | Full data | Exploratory | 35 | $\chi^2(2)=0.09$, p=0.954 | **$\chi^2(1)=9.57$, p=0.002 [β=0.542]** | **$\chi^2(1)=5.57$, p=0.018 [β=0.317]** | $\chi^2(1)=0.09$, p=0.762 [β=-0.080] | $\chi^2(1)=0.03$, p=0.872 [β=0.027] | *$\chi^2(1)=3.00$, p=0.083 [β=0.208]* | $\chi^2(4)=1.56$, p=0.816 |
| | | Base | 35 | | **$\chi^2(1)=11.01$, p<0.001 [β=0.490]** | **$\chi^2(1)=8.34$, p=0.004 [β=0.308]** | — | — | *$\chi^2(1)=3.61$, p=0.057 [β=0.213]* | $\chi^2(4)=1.62$, p=0.806 |
| | Crossing subset | Exploratory | 33 | $\chi^2(3)=2.42$, p=0.491 | **$\chi^2(1)=9.46$, p=0.002 [β=0.449]** | **$\chi^2(1)=4.29$, p=0.038 [β=0.302]** | $\chi^2(1)=0.05$, p=0.826 [β=0.050] | $\chi^2(1)=0.005$, p=0.945 [β=-0.013] | $\chi^2(1)=1.79$, p=0.181 [β=0.196] | $\chi^2(4)=1.99$, p=0.737 |
| | | Base | 33 | | $\chi^2(1)=10.52$, p=0.001 [β=0.486] | $\chi^2(1)=3.02$, p=0.082 [β=0.208] | — | — | — | $\chi^2(4)=2.01$, p=0.734 |

Table S2. Summary of exploratory and base mixed-effects models explaining hair hormone concentrations. Exploratory models contained all candidate predictors and their biologically relevant interaction (competitive status × darting period), whereas base models retained predictors following model simplification. The fifth column reports the likelihood-ratio test comparing each exploratory model to its corresponding base model. Results are presented as Type II Wald $\chi^2$ statistics, degrees of freedom, p-values, and standardised effect sizes (β). Models included a random intercept for individual identity. The crossing subset refers to the base sample of males for which crossing behaviour data were available. Significant effects ($p < 0.05$) are shown in bold and trends in cursive ($p < 0.1$).

| | Service | n | Service vs Base | Service effect | Competitive status | Darting Period | zCSI | Group |
|---|---|---|---|---|---|---|---|---|
| Testosterone | Alarm | 35 | $\chi^2(1)=0.69$, p=0.405 | $\chi^2(1)=0.56$, p=0.453 [β=0.071] | **$\chi^2(1)=18.70$, p<0.001 [β=0.376]** | **$\chi^2(1)=95.12$, p<0.001 [β=0.775]** | — | $\chi^2(4)=4.08$, p=0.395 [β range: -0.243 to -0.057] |
| | BGC | 35 | *$\chi^2(1)=3.47$, p=0.062* | *$\chi^2(1)=2.78$, p=0.095 [β=-0.161]* | **$\chi^2(1)=24.22$, p<0.001 [β=0.416]** | **$\chi^2(1)=58.84$, p<0.001 [β=0.678]** | — | *$\chi^2(4)=8.20$, p=0.084 [β range: -0.343 to -0.091]* |
| | Sentinelling | 35 | $\chi^2(1)=0.30$, p=0.584 | $\chi^2(1)=0.29$, p=0.591 [β=-0.052] | **$\chi^2(1)=18.69$, p<0.001 [β=0.378]** | **$\chi^2(1)=82.43$, p<0.001 [β=0.681]** | — | $\chi^2(4)=3.90$, p=0.420 [β range: -0.347 to -0.094] |
| | Crossing | 33 | *$\chi^2(1)=3.48$, p=0.062* | **$\chi^2(1)=4.73$, p=0.030 [β=-0.195]** | **$\chi^2(1)=17.98$, p<0.001 [β=0.371]** | **$\chi^2(1)=126.30$, p<0.001 [β=0.845]** | — | $\chi^2(4)=3.68$, p=0.452 [β range: -0.328 to -0.100] |
| Cortisol | Alarm | 35 | $\chi^2(1)=0.04$, p=0.851 | $\chi^2(1)=0.02$, p=0.895 [β=-0.020] | **$\chi^2(1)=10.53$, p=0.001 [β=0.491]** | **$\chi^2(1)=5.77$, p=0.016 [β=0.299]** | *$\chi^2(1)=2.75$, p=0.097 [β=0.206]* | $\chi^2(4)=1.59$, p=0.810 [β range: -0.224 to 0.009] |
| | BGC | 35 | **$\chi^2(1)=5.51$, p=0.018** | **$\chi^2(1)=5.43$, p=0.02 [β=-0.430]** | **$\chi^2(1)=14.93$, p<0.001 [β=0.573]** | $\chi^2(1)=0.18$, p=0.669 [β=0.064] | $\chi^2(1)=0.32$, p=0.566 [β=0.077] | $\chi^2(4)=3.99$, p=0.407 [β range: -0.430 to 0.084] |
| | Sentinelling | 35 | $\chi^2(1)=0.22$, p=0.639 | $\chi^2(1)=0.14$, p=0.710 [β=-0.063] | **$\chi^2(1)=10.50$, p=0.001 [β=0.485]** | **$\chi^2(1)=8.20$, p=0.004 [β=0.326]** | *$\chi^2(1)=3.67$, p=0.055 [β=0.213]* | $\chi^2(4)=1.11$, p=0.893 [β range: -0.186 to 0.022] |
| | Crossing | 33 | $\chi^2(1)=0.76$, p=0.382 | $\chi^2(1)=0.67$, p=0.415 [β=-0.119] | **$\chi^2(1)=10.42$, p=0.001 [β=0.484]** | $\chi^2(1)=2.44$, p=0.118 [β=0.186] | — | $\chi^2(4)=1.79$, p=0.775 [β range: -0.134 to 0.105] |

Table S3. Results of mixed-effects models examining the relationship between male service behaviours and hair hormone concentrations. Separate models were fitted for alarm calling, participation in BGC, sentinelling, and crossing behaviour. All service models included the focal service variable in addition to the predictors retained in the corresponding base hormone model (Table S2). The fourth column reports the likelihood-ratio test comparing each service model to its corresponding base hormone model. Results are presented as Type II Wald $\chi^2$ statistics, associated p-values, and standardized effect sizes (β). Models included a random intercept for individual identity. The crossing models were fitted on a base subset of males for which crossing behaviour data were available. Significant effects ($p < 0.05$) are shown in bold and trends in cursive ($p < 0.1$).

RANK MODELS – provided for transparency only

These models are provided for transparency and comparison with the competitive-status models. They are not the models reported or discussed in the main text.

| | | Model | n | Exploratory vs Base | Rank | Darting Period | Rank × Season | Tenure | zCSI | Darting Group |
|---|---|---|---|---|---|---|---|---|---|---|
| Testosterone | Full data | Exploratory | 35 | $\chi^2(3)=2.99$, p=0.393 | **$\chi^2(1)=8.99$, p=0.003 [β=0.203]** | **$\chi^2(1)=63.13$, p<0.001 [β=0.710]** | $\chi^2(1)=0.60$, p=0.440 [β=0.121] | $\chi^2(1)=0.16$, p=0.686 [β=-0.045] | $\chi^2(1)=1.83$, p=0.176 [β=-0.132] | $\chi^2(4)=4.28$, p=0.370 |
| Testosterone | Full data | Base | 35 | $\chi^2(3)=2.99$, p=0.393 | **$\chi^2(1)=8.39$, p=0.004 [β=0.254]** | **$\chi^2(1)=89.49$, p<0.001 [β=0.771]** | — | — | — | $\chi^2(4)=6.22$, p=0.184 |
| Testosterone | Crossing subset | Exploratory | 31 | $\chi^2(3)=3.24$, p=0.357 | **$\chi^2(1)=8.22$, p=0.004 [β=0.192]** | **$\chi^2(1)=73.45$, p<0.001 [β=0.805]** | $\chi^2(1)=1.25$, p=0.264 [β=0.179] | $\chi^2(1)=0.32$, p=0.573 [β=-0.069] | $\chi^2(1)=0.82$, p=0.364 [β=-0.089] | $\chi^2(3)=3.10$, p=0.376 |
| Testosterone | Crossing subset | Base | 31 | $\chi^2(3)=3.24$, p=0.357 | **$\chi^2(1)=10.44$, p=0.001 [β=0.288]** | **$\chi^2(1)=106.41$, p<0.001 [β=0.858]** | — | — | — | $\chi^2(3)=2.37$, p=0.499 |
| Cortisol | Full data | Exploratory | 35 | $\chi^2(2)=3.10$, p=0.213 | **$\chi^2(1)=11.44$, p<0.001 [β=0.133]** | **$\chi^2(1)=6.60$, p=0.010 [β=0.288]** | $\chi^2(1)=1.93$, p=0.164 [β=0.299] | $\chi^2(1)=0.05$, p=0.815 [β=-0.041] | *$\chi^2(1)=2.85$, p=0.091 [β=0.167]* | $\chi^2(4)=2.39$, p=0.664 |
| Cortisol | Full data | Base | 35 | $\chi^2(2)=3.10$, p=0.213 | **$\chi^2(1)=9.81$, p=0.002 [β=0.373]** | **$\chi^2(1)=7.99$, p=0.005 [β=0.313]** | — | — | $\chi^2(1)=1.69$, p=0.194 [β=0.157] | $\chi^2(4)=2.50$, p=0.644 |
| Cortisol | Crossing subset | Exploratory | 31 | **$\chi^2(3)=22.74$, p<0.001** | **$\chi^2(1)=9.28$, p=0.002 [β=0.081]** | **$\chi^2(1)=6.81$, p=0.009 [β=0.295]** | $\chi^2(1)=2.30$, p=0.130 [β=0.355] | $\chi^2(1)=0.07$, p=0.798 [β=-0.049] | $\chi^2(1)=1.69$, p=0.193 [β=0.151] | $\chi^2(3)=1.92$, p=0.590 |
| Cortisol | Crossing subset | Base | 31 | **$\chi^2(3)=22.74$, p<0.001** | **$\chi^2(1)=6.90$, p=0.009** | *$\chi^2(1)=3.81$, p=0.051* | — | — | — | $\chi^2(3)=2.34$, p=0.505 |

Table S4. Summary of exploratory and base mixed-effects models explaining hair hormone concentrations but with rank instead. Exploratory models contained all candidate predictors and their biologically relevant interaction (rank × darting period), whereas base models retained predictors following model simplification. The fifth column reports the likelihood-ratio test comparing each exploratory model to its corresponding base model. Results are presented as Type II Wald $\chi^2$ statistics, degrees of freedom, p-values, and standardised effect sizes (β). Models included a random intercept for individual identity. The crossing subset refers to the base sample of males for which crossing behaviour data were available. Significant effects ($p < 0.05$) are shown in bold and trends in cursive ($p < 0.1$).

| | Service | n | Service vs Base | Service effect | Rank | Darting Period | zCSI | Group |
|---|---|---|---|---|---|---|---|---|
| Testosterone | Alarm | 35 | $\chi^2(1)=0.25$, p=0.618 | $\chi^2(1)=0.25$, p=0.621 [β=0.053] | **$\chi^2(1)=7.74$, p=0.005 [β=0.247]** | **$\chi^2(1)=81.53$, p<0.001 [β=0.785]** | — | $\chi^2(4)=4.25$, p=0.374 [β range: -0.366 to -0.129] |
| | BGC | 35 | $\chi^2(1)=0.53$, p=0.468 | $\chi^2(1)=0.35$, p=0.555 [β=-0.068] | **$\chi^2(1)=8.53$, p=0.004 [β=0.264]** | **$\chi^2(1)=56.26$, p<0.001 [β=0.741]** | — | $\chi^2(4)=6.46$, p=0.167 [β range: -0.429 to -0.160] |
| | Sentinelling | 35 | $\chi^2(1)=0.61$, p=0.433 | $\chi^2(1)=0.50$, p=0.480 [β=-0.078] | **$\chi^2(1)=8.41$, p=0.004 [β=0.256]** | **$\chi^2(1)=75.11$, p<0.001 [β=0.801]** | — | $\chi^2(4)=4.53$, p=0.340 [β range: -0.371 to -0.132] |
| | Crossing | 31 | **$\chi^2(1)=9.69$, p=0.002** | **$\chi^2(1)=15.31$, p<0.001 [β=-0.277]** | **$\chi^2(1)=23.48$, p<0.001 [β=0.315]** | **$\chi^2(1)=218.32$, p<0.001 [β=0.828]** | — | $\chi^2(3)=4.16$, p=0.245 [β range: 0.063 to 0.204] |
| Cortisol | Alarm | 35 | $\chi^2(1)=0.00$, p=0.971 | $\chi^2(1)=0.01$, p=0.931 [β=-0.014] | **$\chi^2(1)=9.31$, p=0.002 [β=0.377]** | **$\chi^2(1)=5.57$, p=0.018 [β=0.305]** | $\chi^2(1)=1.24$, p=0.266 [β=0.149] | $\chi^2(4)=2.38$, p=0.665 [β range: -0.319 to 0.006] |
| | BGC | 35 | **$\chi^2(1)=3.90$, p=0.048** | **$\chi^2(1)=4.29$, p=0.038 [β=-0.386]** | **$\chi^2(1)=12.19$, p<0.001 [β=0.460]** | $\chi^2(1)=0.28$, p=0.594 [β=0.084] | $\chi^2(1)=0.00$, p=0.977 [β=0.004] | $\chi^2(4)=4.56$, p=0.335 [β range: -0.377 to 0.048] |
| | Sentinelling | 35 | $\chi^2(1)=0.09$, p=0.760 | $\chi^2(1)=0.04$, p=0.834 [β=-0.036] | **$\chi^2(1)=9.27$, p=0.002 [β=0.368]** | **$\chi^2(1)=7.18$, p=0.007 [β=0.324]** | $\chi^2(1)=1.66$, p=0.197 [β=0.158] | $\chi^2(4)=1.97$, p=0.741 [β range: -0.300 to 0.008] |

| | | | | | | | | |
|---|---|---|---|---|---|---|---|---|
| | Crossing | 31 | $\chi^2(1)$=0.86, p=0.354 | $\chi^2(1)$=0.67, p=0.414 [β=-0.128] | **$\chi^2(1)$=7.03, p=0.008 [β=0.380]** | *$\chi^2(1)$=3.24, p=0.072 [β=0.230]* | — | $\chi^2(3)$=2.57, p=0.462 [β range: 0.081 to 0.254] |

Table S5. Results of mixed-effects models examining the relationship between male service behaviours and hair hormone concentrations. Separate models were fitted for alarm calling, participation in BGC, sentinelling, and crossing behaviour. All service models included the focal service variable in addition to the predictors retained in the corresponding base hormone model (Table S2). The fourth column reports the likelihood-ratio test comparing each service model to its corresponding base hormone model. Results are presented as Type II Wald $\chi^2$ statistics, associated p-values, and standardized effect sizes (β). Models included a random intercept for individual identity. The crossing models were fitted on a base subset of males for which crossing behaviour data were available. Significant effects ($p < 0.05$) are shown in bold and trends in cursive ($p < 0.1$).
